# Systematic Evaluation of Transfer Learning Strategies for Clinical Chemotherapy Response Prediction

**DOI:** 10.64898/2026.02.16.706121

**Authors:** Hanqin Du, Pedro Ballester

## Abstract

Accurately predicting chemotherapy response remains a major challenge in precision oncology. Although machine-learning models based on tumour omics data have shown promise, the majority of existing studies are trained and evaluated on pre-clinical cell-line datasets, leaving their clinical applicability insufficiently characterised. In this study, we systematically evaluate a range of transfer-learning strategies for chemotherapy response prediction under realistic clinical constraints using patient data from The Cancer Genome Atlas (TCGA). Rather than proposing a new predictive model, we focus on assessing the effectiveness and limitations of commonly used approaches for transferring pre-clinical knowledge to clinical settings. These include cell-line-validated biomarkers, biologically informed feature representations, direct application of pre-clinical deep-learning models, model fine-tuning, and hybrid strategies that integrate pre-clinical predictions with clinical data. All methods are evaluated within a unified framework using consistent cohort construction, shared performance metrics, and bias-controlled validation procedures. Across multiple drugs and molecular data types, we find that most transfer strategies—including biomarker-based feature selection and direct pre-clinical model transfer—fail to produce robust or consistent improvements in clinical prediction performance. In contrast, conservative approaches based on fine-tuning pre-clinical models or incorporating pre-clinical predictions as features in clinical models yield more stable and reproducible gains. Further improvements are observed when basic pre-treatment clinical variables are integrated. Together, our results demonstrate the practical boundaries of pre-clinical to clinical transfer for drug response prediction and highlight hybrid and fine-tuning strategies as more reliable baselines for future translational modelling efforts.

## Introduction

Chemotherapy is one of the most widely used cancer treatments due to its broadly applicability, affordability, and routinely administered either alone or in combination with surgery, radiotherapy, targeted therapy, and immunotherapy^1^. However, chemotherapy effectiveness varies between patients and tumour types. The response rates can be high in some contexts—for example, anthracycline-based regimens in acute promyelocytic leukaemia improve further when combined with all-trans retinoic acid and arsenic trioxide^2^—yet remain modest in others, such as gemcitabine monotherapy in pancreatic cancer (∼7%)^3^. Since chemotherapy can carry significant toxicity and delay access to more effective alternatives, accurally predicting response prior to treatment is of importance for improving outcomes and reducing unnecessary burden for patients.

This variability is closely linked to the multifactorial nature of drug resistance. Cancer cells may reduce drug uptake, increase efflux, alter drug metabolism, evade apoptosis, or modify drug targets, and multiple mechanisms can coexist within a single tumour^4^. In clinical practice, predictive biomarkers are used where available, but robust, guideline-endorsed biomarkers exist only for a minority of chemotherapy settings. Moreover, many biomarkers operate as threshold tests (“positive/negative”), which limits their usefulness when multiple therapies are available^5^. Even widely used examples illustrate this limitation: MGMT promoter methylation is an established predictor of temozolomide sensitivity, yet treatment response is also shaped by other DNA repair pathways and broader cellular programmes^6^. These realities motivate approaches that integrate information across different molecular features rather than relying on a single marker.

Machine learning (ML) provides a solution for building multivariate predictors from tumour omics profiles. Transcriptomic, epigenetic, and genomic measurements can capture aspects of cellular state relevant to drug response and can be combined to model interactions. However, clinical translation is constrained by the nature of real-world patient data. Compared with pre-clinical resources, clinical cohorts with matched omics profiles and treatment response labels are typically small, heterogeneous, and mostly imbalanced. In TCGA, drug-treated subsets vary widely across drugs and tumour types, often yielding only dozens of labelled samples per drug. This creates the classical “small data” regime (*p* ≫ *n*), where models are prone to overfitting and reported performance is highly sensitive to evaluation design^7,8^. These issues are exacerbated by label noise and clinical confounding: clinical response is influenced not only by tumour resistance mechanisms but also by patient-level factors such as health status, comorbidities, tumour grade, and treatment context. Consequently, evaluation must rely on metrics suited to imbalance and small samples, such as MCC and ROC–AUC, and must control for optimistic bias introduced by repeated model selection and unstable splits^9,10^.

Because of these limitations, the literature on drug-response prediction has been dominated by large-scale pre-clinical pharmacogenomic datasets, especially GDSC and CCLE^11,12,13^. These resources provide thousands of cell lines screened against hundreds of compounds, typically with response quantified as IC50, enabling the training of complex models—often deep neural networks—and the use of regression metrics such as RMSE and Pearson correlation^14^. Methodological innovations have proliferated in this setting, including multi-view and multi-omics embedding approaches^15^, similarity-network models that exploit relationships between cell lines and drugs^16^, and autoencoder-based representation learning to address high-dimensional inputs across molecular layers^17,18,19^. Graph neural networks have also been used to incorporate biological network priors, such as co-expression or protein–protein interaction structure, and to integrate drug molecular representations^20,21,22^. Though these developments demonstrate strong predictive performance in vitro, the translational value of cell-line-trained models is debated since cell lines differ from patient tumours in critical ways, including the absence of tumour microenvironment, altered baseline gene expression, and reduced heterogeneity^23,24^. As a result, models trained exclusively on pre-clinical data often show a notable performance drop when applied to patient tumours, and systematic clinical validation remains relatively uncommon^14^.

Transfer learning has therefore been proposed as a principled approach to leverage data-rich pre-clinical resources for data-poor clinical prediction^25,26^. In this domain, “transfer learning” should be interpreted broadly. Beyond the familiar pre-training and fine-tuning paradigm, transfer can occur via feature-level reuse (e.g., projecting gene expression into pathway activities), parameter transfer (fine-tuning parts of a pre-trained model), instance re-weighting, explicit domain alignment, or hybrid strategies that combine pre-clinical and clinical signals. Knowledge-driven feature extraction methods—such as PROGENy^27^, GSVA using MSigDB Hallmark gene sets^28,29^, and DoRothEA^30^—map transcriptomes into biologically interpretable activities and may reduce dimensionality and noise, potentially improving robustness under small clinical sample sizes. Model-level transfer from cell lines to tumours has also been explored: MOLI is one of the few deep-learning frameworks that explicitly incorporates batch-effect correction and reports clinical evaluation as part of its pre-clinical-to-clinical pipeline^31^. Nevertheless, despite the popularity of transfer-learning concepts in this area, their practical effectiveness, stability, and limitations under realistic clinical constraints are still insufficiently characterised in a unified and bias-controlled setting.

In this study, we present a systematic and bias-controlled evaluation of transfer-learning strategies for chemotherapy response prediction under clinical constraints. Rather than introducing a new predictive architecture, we benchmark multiple representative approaches—including literature-derived cell-line biomarkers, biologically informed feature representations, direct transfer of pre-clinical deep-learning models, fine-tuning, and hybrid transfer—within a unified evaluation framework. All methods are assessed using consistently constructed TCGA cohorts, shared performance metrics, and nested cross-validation with explicit bias correction. Our results demonstrate that most pre-clinical transfer strategies fail to deliver robust gains in clinical settings, whereas conservative fine-tuning and hybrid approaches yield more stable improvements. These findings highlight the importance of rigorous evaluation design and provide a reproducible reference for future studies on cross-domain drug response prediction.

## Methods

### Data sources and cohort construction

All clinical and molecular data analysed in this study were obtained from The Cancer Genome Atlas (TCGA) and downloaded from the Genomic Data Commons (GDC) repository. Patients who received chemotherapy before sample collection were excluded, as such exposure may induce treatment resistance and bias downstream prediction. For patients with multiple molecular profiles available, only a single sample was retained to avoid repeated measurements from the same individual. Specifically, the molecular profile corresponding to the earliest available sampling time point relative to tumour diagnosis was selected. When multiple samples were available at the same time point, the first sample was chosen based on alphanumeric ordering of TCGA sample identifiers.

Pre-clinical drug response data were obtained from the Genomics of Drug Sensitivity in Cancer (GDSC) project. Continuous IC50 measurements and corresponding binarised drug response labels were retrieved from Iorio *et al.*^12^, wehre drug response binarisation was performed using the LOBICO framework (Logic Optimization for Binary Input to Continuous Output)^32^.

To reduce systematic distributional differences between pre-clinical and clinical transcriptomic data, ComBat batch-effect correction was applied to align TCGA samples with GDSC data prior to model transfer. For machine-learning algorithms sensitive to feature scaling—including support vector machines, linear models, and deep learning approaches—mRNA expression values were converted to transcripts per million (TPM) before downstream analysis. To ensure a fair and representative evaluation, we focused on chemotherapeutic agents that are widely used across multiple cancer types and sufficiently represented in both TCGA and GDSC. Based on these criteria, four drugs—cisplatin, fluorouracil, gemcitabine, and paclitaxel—were selected for subsequent analyses; detailed selection criteria are provided in Appendix.

### Molecular and biomarker features

For each of the four evaluated chemotherapeutic agents, we curated a set of candidate drug-resistance biomarkers from the published literature. Biomarkers were included only if the corresponding studies reported experimental validation in pre-clinical systems, primarily in vitro cancer cell-line models. Where available, evidence from more clinically relevant systems—such as cell line–derived xenografts (CDX) and patient-derived xenografts (PDX)—was also incorporated. In total, the curated biomarker sets comprised 81 candidates for cisplatin, 53 for fluorouracil, 44 for gemcitabine, and 55 for paclitaxel. Due to the size of these collections, the complete biomarker lists and their supporting references are provided in the Appendix.

To construct biomarker-based feature matrices, we adopted an inclusive strategy that aimed to retain as much prior biological information as possible. Specifically, when a publication reported a biomarker at the gene or miRNA family level (e.g. *miR-7*), all corresponding molecular features available in the omics data (e.g. *miR-7-1* and *miR-7-2*) were retained. This approach avoids prematurely excluding potentially relevant molecular signals and ensures comprehensive coverage of reported biomarkers.

These biomarker-derived features were used as a form of prior-knowledge–based feature selection rather than as ground-truth clinical predictors. Their purpose was to assess whether restricting model inputs to experimentally supported resistance-related features could improve predictive performance or stability relative to models trained on raw omics profiles. All biomarker features were extracted from the same preprocessed omics matrices used for the corresponding raw-omics models, ensuring that any performance differences reflect feature selection rather than differences in data processing.

### Pathway activation inference

To incorporate prior biological knowledge into downstream machine-learning models, bulk mRNA expression profiles were transformed into biologically informed activity scores capturing signalling pathways, biological processes, and transcription factor (TF) regulation.

Signalling pathway activities were inferred using PROGENy^27^, a perturbation-based framework that estimates pathway activity as a weighted linear combination of responsive genes derived from large-scale experimental perturbation data. Using the *progeny* R package (v3.22), activity scores were computed for 14 canonical signalling pathways. Biological process activities were estimated with Gene Set Variation Analysis (GSVA)^33^, a non-parametric enrichment method applied on a per-sample basis. GSVA was performed using the 50 Hallmark gene sets from MSigDB, which summarise core biological processes while reducing redundancy. Transcription factor activities were inferred using DoRothEA^30^ via the *decoupleR* package (v1.22.0), leveraging curated TF– target regulons. Only interactions with confidence levels A–C were retained.

Together, these transformations reduced the transcriptomic feature space to 236 biologically interpretable variables, comprising 14 pathway activities, 50 Hallmark scores, and 172 TF activities. These representations provide structured, low-dimensional inputs for downstream modelling and enable assessment of biologically informed feature transfer under small-sample clinical conditions.

### Traditional machine-learning evaluation with nested cross-validation

Given the limited sample size of clinical cohorts, we focused on traditional machine-learning (TRAD) models that are known to be more robust under small-*n*, high-*p* conditions. To systematically evaluate the predictive potential of biomarker-based and biologically inferred features, we adopted a conservative evaluation framework previously described in our earlier work^34^.

Briefly, six widely used machine-learning algorithms were considered: logistic regression, support vector machines (SVM), classification and regression trees (CART), random forests (RF), XGBoost, and LightGBM. Model performance was evaluated using nested cross-validation with 10 outer folds for unbiased performance estimation and 5 inner folds for hyperparameter tuning. Optimal Model Complexity (OMC)–based feature selection was included as a tunable hyperparameter.

To account for randomness in data splitting and model initialisation, all evaluations were repeated with five different random seeds, and median performance across repetitions was reported. Both threshold-dependent and threshold-independent metrics were used, specifically Matthews correlation coefficient (MCC) and ROC–AUC. Bootstrap bias correction (BBC)^35^ was applied to estimate the performance of the best-selected model while correcting for overestimation induced by evaluating multiple model on the same dataset.

### Fine-tuning of the pretrained MOLI model on TCGA data

To adapt pre-clinical deep-learning models to patient-level drug response prediction, we performed supervised fine-tuning of pretrained MOLI models using TCGA gene expression data. Models were initialised from checkpoints trained on GDSC, including network weights and feature normalisation parameters. Fine-tuning was evaluated using stratified five-fold cross-validation on TCGA. In each fold, one subset was held out for testing, while the remaining samples were used for training and validation with early stopping. Given the limited size of clinical cohorts, a conservative strategy was adopted in which the pretrained encoder was frozen and only the classifier layers was updated during fine-tuning. Standard supervised optimisation settings were used, with class imbalance handled via weighted sampling. Out-of-fold predictions from all folds were aggregated to obtain a complete set of unbiased predictions across the cohort. Predictive performance was quantified using ROC–AUC computed on these aggregated predictions, providing a robust estimate of generalisation performance under small-sample clinical conditions.

### Hybrid transfer strategy

To integrate information learned from pre-clinical data into clinical drug-response prediction, we adopted a conservative prediction-level transfer strategy (**Figure 1**). A deep-learning model (MOLI) was first trained on GDSC cell-line data to predict drug response. For each TCGA patient sample, the output of the pretrained model was computed and used as an additional continuous feature, referred to as the *cell-line score*.

**Figure 1.**
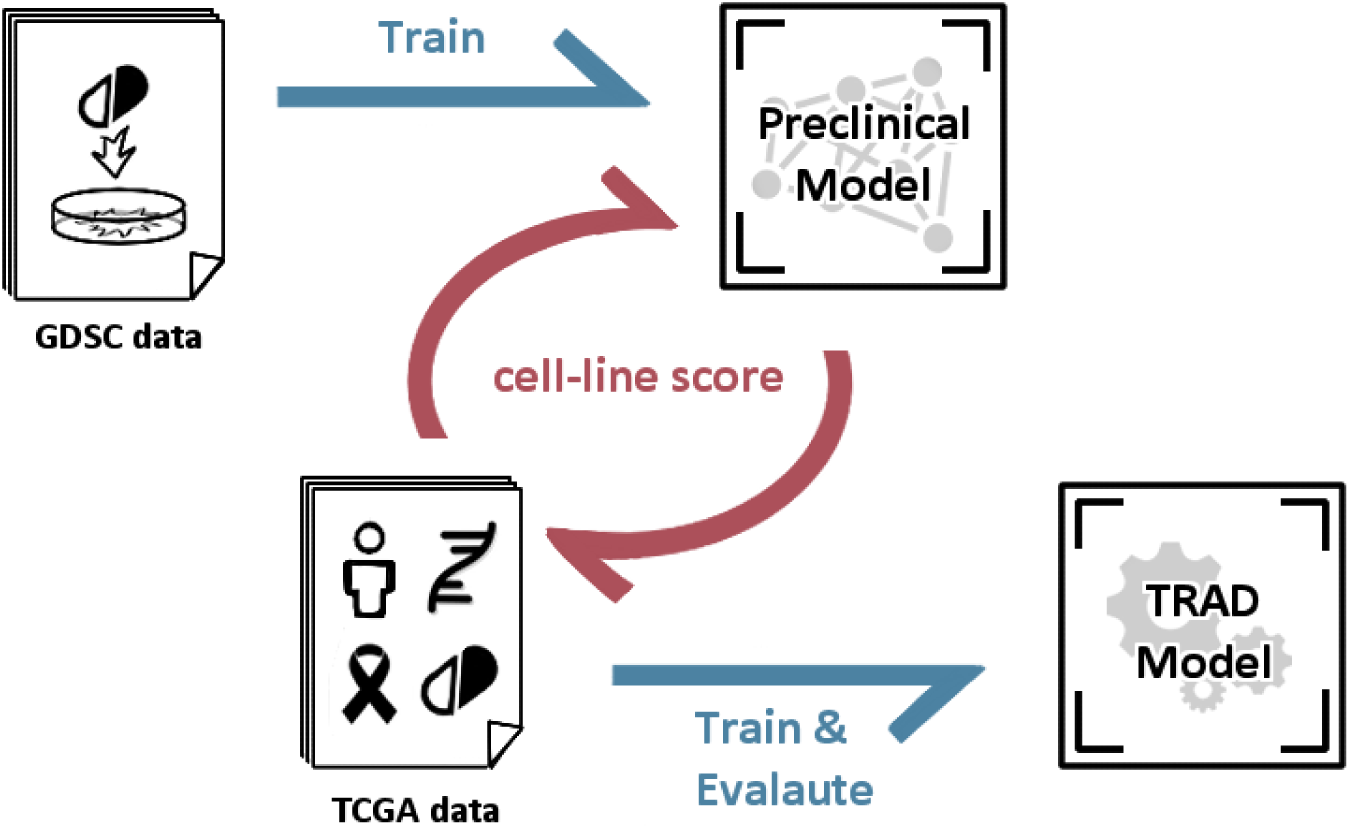
Overview of the hybrid transfer-learning strategy. A deep-learning model is first trained on pre-clinical cell-line data (GDSC) to predict drug response. The resulting prediction score for each sample, referred to as the cell-line score, is then transferred to the clinical domain and used as an additional input feature alongside clinical omics data. A conventional machine-learning (TRAD) model is trained and evaluated on TCGA samples using this augmented feature set. The pre-clinical model remains fixed during clinical model training, enabling prediction-level transfer without parameter updating or domain re-training.

The cell-line score was concatenated with clinical omics features, and standard machine-learning classifiers were trained and evaluated on TCGA data using the same nested cross-validation and bias-correction framework described above. Importantly, no parameters of the pre-clinical model were updated during this process, avoiding assumptions of full domain alignment. Model performance was assessed using ROC–AUC, and bootstrap bias correction was applied when comparing multiple models to control for selection-induced optimism.

### Clinical features extraction and integration

To assess whether basic clinical information provides complementary predictive value in the hybrid transfer setting, a small set of pre-treatment clinical variables was incorporated into the models. Specifically, patient age, time interval between biospecimen procurement and therapy initiation, tumour grade, specimen procurement type, and Karnofsky performance score were extracted from TCGA clinical records.

## Results

### Pre-clinical and clincial dataset composition

We used clinical drug-response data from The Cancer Genome Atlas (TCGA) to evaluate transfer-learning strategies from pre-clinical to clinical settings. To ensure that the selected drugs were suitable for cross-domain evaluation, we focused on chemotherapeutic agents that (i) were present in both TCGA and GDSC, (ii) had relatively large sample sizes in clinical cohorts. Based on these criteria, four widely used drugs—Cisplatin, Fluorouracil, Gemcitabine, and Paclitaxel—were selected for subsequent analyses.

**Table 1** summarises the sample composition of the pre-clinical training sets (GDSC) and the corresponding clinical evaluation cohorts (TCGA). In GDSC, drug response is binarised from IC50 values, whereas in TCGA, response labels is binarised from Recist-based drug response classifications and are dichotomised as responder (R) or non-responder (NR). Across all four drugs, the pre-clinical datasets are substantially larger than the clinical cohorts and exhibit markedly different response distributions, highlighting the domain shift between cell-line and patient data. **Figure 2** illustrates the response-label distribution for Cisplatin after this integration step. Equivalent distributions for Fluorouracil, Gemcitabine, and Paclitaxel are provided in the Appendix.

**Figure 2.**
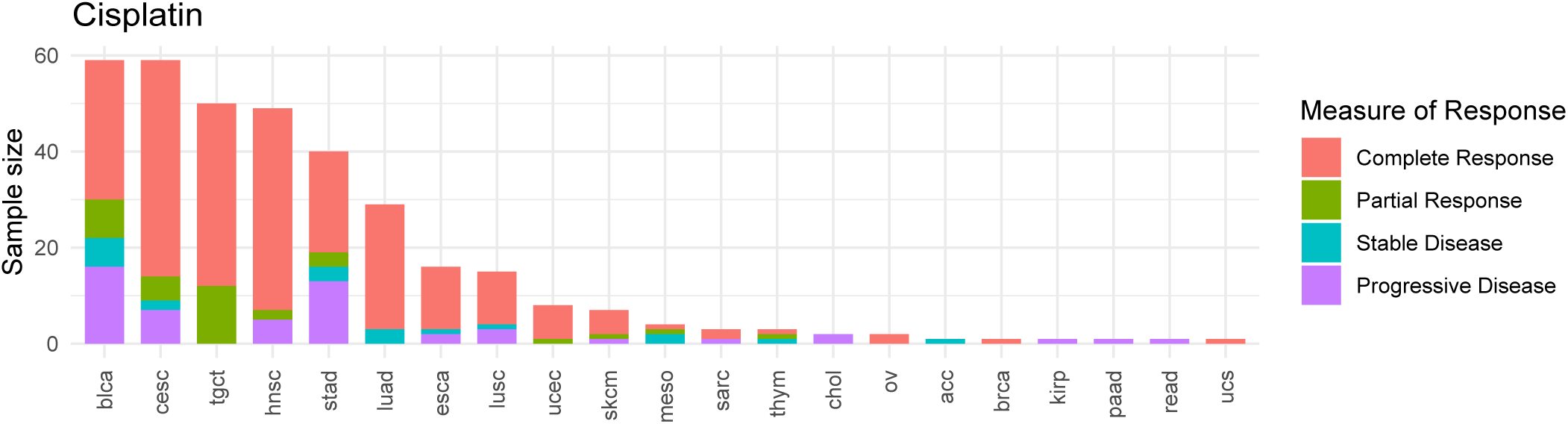
Distribution of responder (R) and non-responder (NR) samples across cancer types for Cisplatin in the TCGA cohort, after merging low-sample or highly imbalanced tumour types into the *Other* category..

**Table 1.** Summary of drug-response sample composition in the GDSC training sets and TCGA clinical cohorts. R and NR represent responder and non-responder samples, respectively.

| Variable | GDSC train set | TCGA test set |
| --- | --- | --- |
| Cisplatin | 829 (R 77, NR 752) | 352 (R 279, NR 73) |
| Fluorouracil | 889 (R 92, NR 797) | 245 (R 179, NR 66) |
| Gemcitabine | 844 (R 54, NR 790) | 170 (R 81, NR 89) |
| Paclitaxel | 389 (R 26, NR 363) | 204 (R 147, NR 57) |

### Limited predictive value of pre-clinical biomarkers in clinical drug response prediction

We next evaluated whether biomarkers validated in pre-clinical cell-line experiments could improve drug response prediction in clinical cohorts. For each of the four selected chemotherapeutic agents—Cisplatin, Fluorouracil, Gemcitabine, and Paclitaxel—we selected a comprehensive set of drug-resistance–associated biomarkers from the literature. Each biomarker had been experimentally validated in cell-line *in vivo* or xenograft models, and for each drug more than 50 candidate biomarkers were collected. A complete list of biomarkers and their corresponding references is provided in the Appendix.

To assess the predictive value of these biomarkers in a clinical setting, we trained drug-specific pan-cancer models on TCGA data using either raw omics features or biomarker-restricted feature sets. Model training and evaluation followed the same nested cross-validation framework described above, using six commonly applied machine-learning algorithms. For consistency with subsequent analyses, models were evaluated under a mixed-cancer setting, and predictive performance was quantified using both Matthews correlation coefficient (MCC) and ROC–AUC.

**Figure 3** summarises the performance distributions across drugs and feature types. Across all four drugs, models trained on biomarker-restricted feature sets did not exhibit systematically higher predictive performance than models trained on the corresponding raw omics data. In most settings, median MCC and ROC–AUC values were comparable between biomarker-based and omics-based models, with overlapping performance distributions across algorithms. Moreover, biomarker-based models did not demonstrate improved stability, as reflected by similar or wider variability in performance across cross-validation runs.

**Figure 3.**
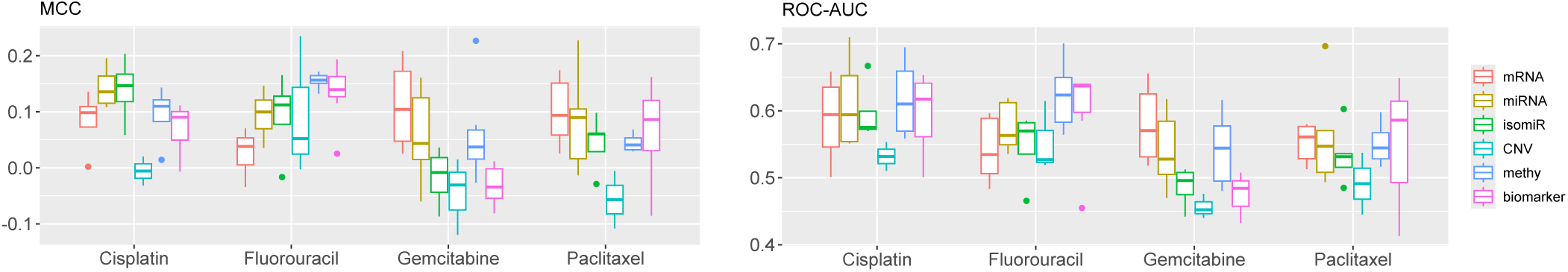
Comparison of predictive performance for mixed-cancer models trained using raw omics features and literature-derived biomarker feature sets. Boxplots show the distribution of median MCC (left) and ROC–AUC (right) values across model runs. Each point represents the median performance of a single algorithm evaluated over five repeated nested cross-validation runs.

These results indicate that restricting model inputs to biomarkers derived from pre-clinical experiments does not, by itself, enhance predictive accuracy or robustness in clinical drug response prediction. Cancer-type–specific comparisons and detailed results for individual tumour groups are provided in the Appendix.

### Feature-level transfer using pathway-based feature transfer

To assess whether biologically informed feature representations provide advantages over raw transcriptomic inputs, we compared models trained on mRNA expression profiles with those trained on inferred pathway-level features. Pathway activities were obtained using PROGENy, GSVA Hallmark gene sets, and DoRothEA, yielding a total of 236 biologically derived features. Model training and evaluation followed the unified machine-learning pipeline introduced in previous section. No additional feature selection was applied prior to training to allowing a direct comparison between models using either raw mRNA inputs or inferred pathway activities. Performance was evaluated across multiple drug–cancer-type combinations using the MCC, which provides a balanced measure of predictive accuracy under class imbalance.

Across the evaluated drug–cancer combinations, models trained on pathway-derived features achieved largely similar performance compare to those trained directly on mRNA expression profiles (**Figure 4**). Statistical significance was assessed using two-sided Wilcoxon tests. In drug–cancer pairs where mRNA-based models produced positive predictive performance (e.g., Cisplatin–LUSC, Cisplatin–other), the pathway-based models also achieved positive MCC values. In combinations where mRNA features did not yield predictive performance, pathway-based features similarly did not result in improved MCC. In specific cases, pathway-derived features were associated with higher MCC (e.g., Cisplatin–LUAD, Gemcitabine–other, Fluorouracil–other, Paclitaxel–other). In contrast, lower MCC values were observed for pathway-based models in settings such as Paclitaxel–HNSC.

**Figure 4.**
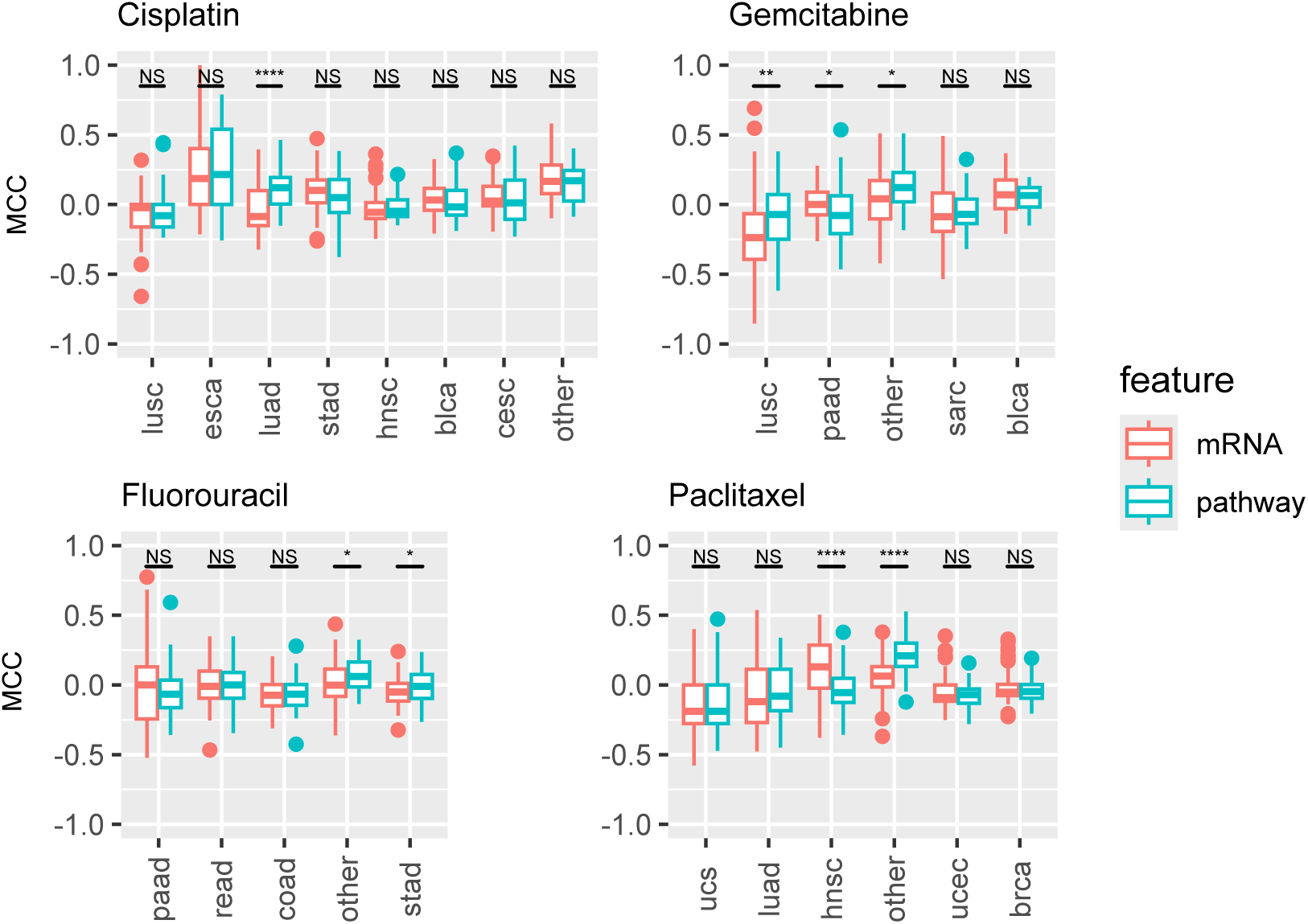
Comparison of model performance using mRNA profiles (red) and inferred pathway activities (blue). Each boxplot shows the distribution of MCC values across model runs. Statistical significance was evaluated using two-sided Wilcoxon tests.

These results indicate that pathway-level abstractions preserve much of the predictive information contained in raw transcriptomic data and substantially reduce feature dimensionality, but do not consistently enhance predictive performance.

### Model-level transfer from pre-clinical deep learning

We next evaluated model-level transfer from pre-clinical deep learning models to clinical drug response prediction using the MOLI framework. MOLI was selected as a representative multi-omics deep-learning model designed for drug response prediction, with publicly available code and preprocessed training set. Following the original study, MOLI models were trained on GDSC cell-line data using mRNA expression profiles as input.

Hyperparameters were selected from 50 randomly sampled configurations using 5-fold cross-validation, and model training employed early stopping based on validation loss. Since 5-fold cross-validation was used during training, five models with partially overlapping training data but different hyperparameter configurations were obtained for each drug. Predictive performance was summarised using the median ROC–AUC across folds, with the minimum and maximum values reported to reflect training variability.

We first evaluated the trained MOLI models by directly applying them to our harmonised TCGA cohorts. Gene expression profiles from GDSC and TCGA were aligned using ComBat batch-effect correction prior to inference. Under this direct transfer setting, the MOLI models achieved moderate predictive performance for Cisplatin and Fluorouracil, with median ROC–AUC values of 0.64 and 0.55, respectively. In contrast, performance for Gemcitabine and Paclitaxel was close to the non-informative baseline, with median ROC–AUC values of 0.51 and 0.43.

Two strategies to adapt the pre-clinical MOLI models to clinical data is explored. First, fine-tuning approach in which the encoder layers of the pre-trained MOLI models were frozen, and the downstream prediction layers were updated using TCGA data. Fine-tuning and evaluation were performed using nested cross-validation, with the inner loop used for training and early stopping and the outer loop reserved for unbiased performance estimation. Second, a conservative hybrid strategy in which predictions generated by the pre-trained MOLI models were used as an additional feature (“cell-line score”) in traditional machine-learning models trained on TCGA data. Six commonly used machine-learning algorithms were evaluated, and predictive performance was summarised using bootstrap bias–corrected (BBC) ROC–AUC values to mitigate overestimation arising from repeated model selection. As shown in **Figure 5**, both fine-tuning and hybrid approach yielded certain degree of improvements over direct application of MOLI and reduced performance variability across all four drugs.

**Figure 5.**
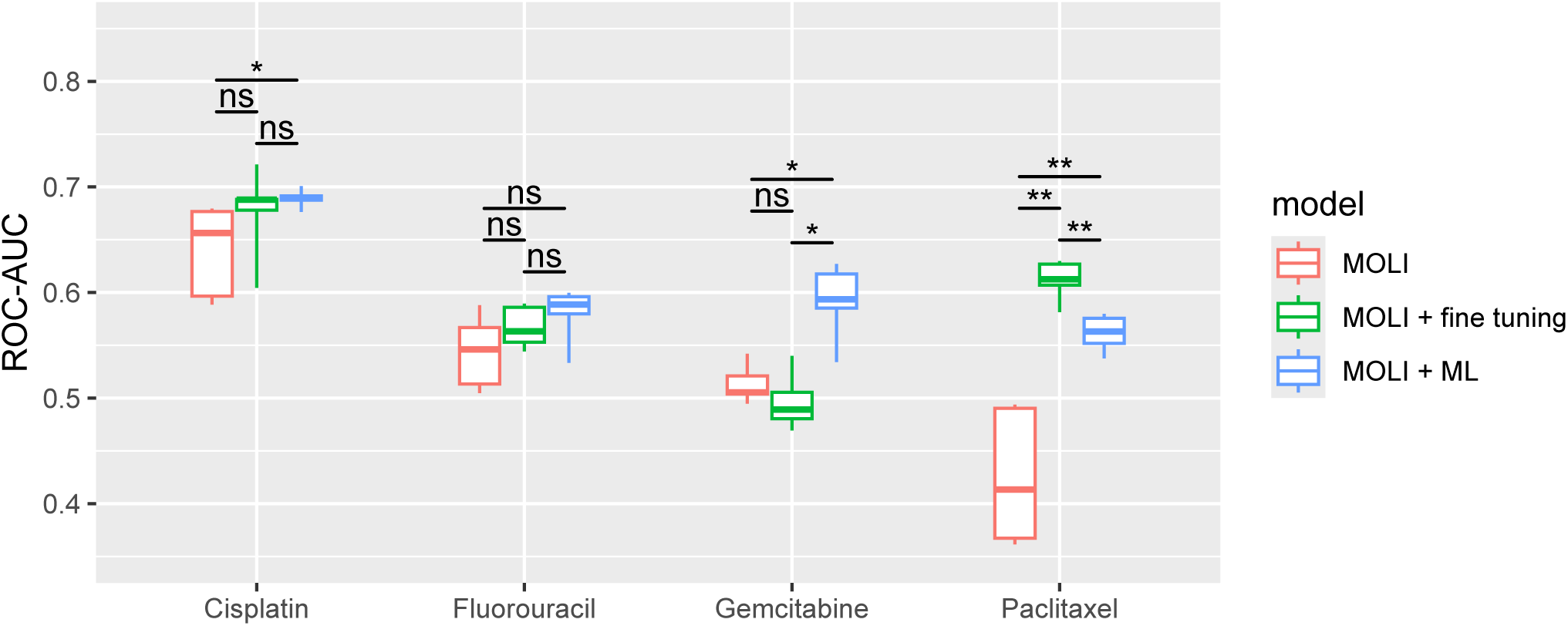
Comparison of ROC–AUC performance for three model-level transfer strategies across four drugs: direct application of pre-trained MOLI models (red), fine-tuned MOLI models (green), and hybrid models combining MOLI predictions with clinical machine-learning models (blue). Boxplots show the distribution of ROC–AUC values across repeated evaluations. Statistical significance was assessed using two-sided Wilcoxon tests.

Together, these results indicate that naïve parameter adaptation of pre-clinical deep-learning models is insufficient to achieve robust clinical generalisation. Instead, integrating pre-clinical model outputs with clinical data in a hybrid framework provides a more stable and effective transfer strategy.

### hybrid transfer allows clinical feature integration

To further assess whether clinical information provides complementary value in the transfer-learning setting, we extended the hybrid models by incorporating four pre-treatment clinical variables: age, Karnofsky performance score, tumour grade, and treatment date. All variables were collected prior to treatment initiation to avoid data leakage. This analysis was performed for Cisplatin, Gemcitabine, and Paclitaxel, where sufficient clinical annotation was available. Model performance was evaluated using bootstrap bias–corrected (BBC) median ROC–AUC values to mitigate optimistic bias arising from repeated model evaluation. For each drug–omics combination, we compared models trained on omics features alone with models additionally incorporating clinical variables (**Figure 6**).

**Figure 6.**
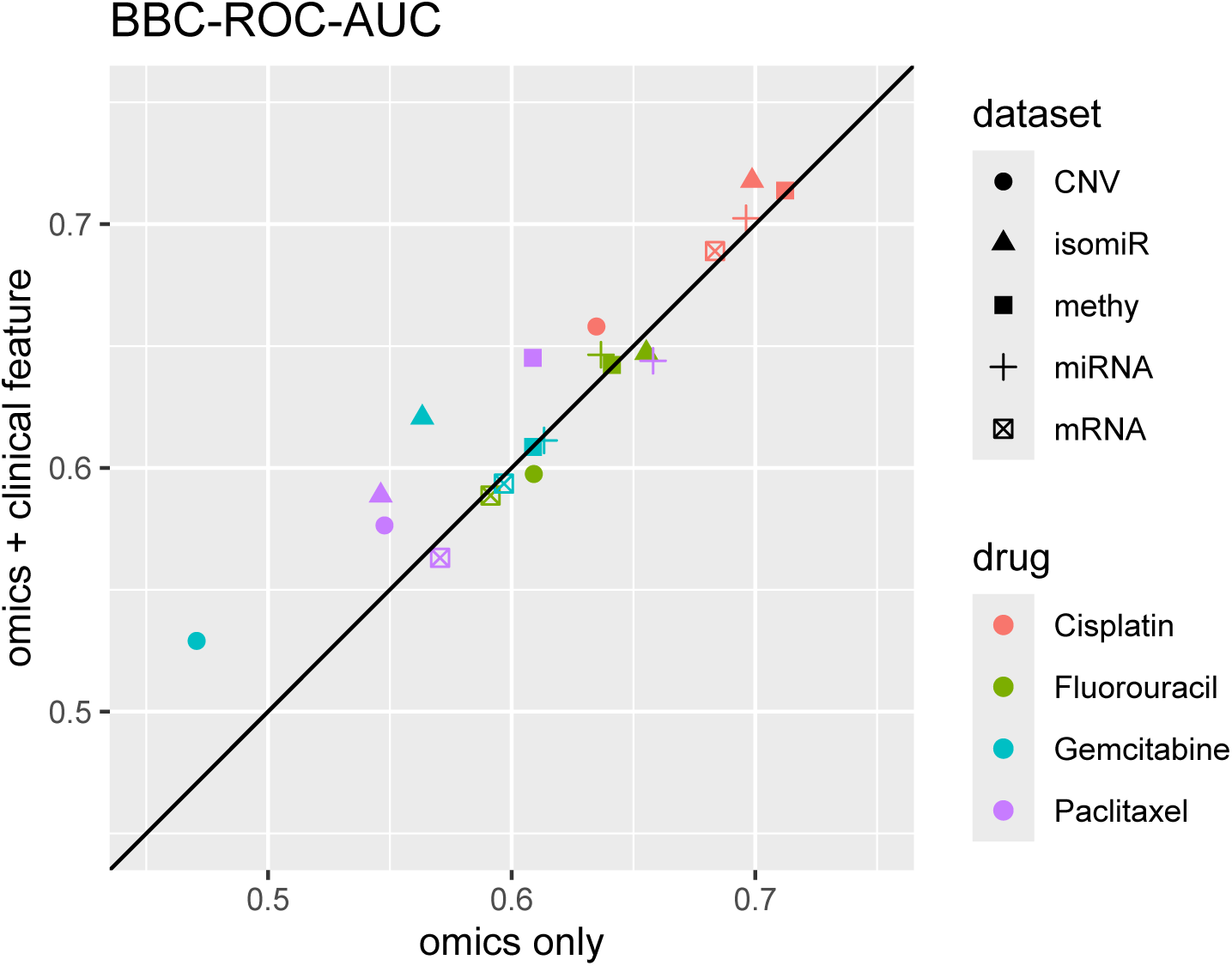
Performance comparison of models trained with and without additional clinical features. Each point corresponds to a specific drug–omics combination. The diagonal line represents equal performance under both configurations.

Across all evaluated settings, the inclusion of clinical features resulted in performance improvements in approximately half of the drug–omics combinations. Improvements were observed consistently across multiple molecular layers, including mRNA, miRNA, DNA methylation, isomiR, and CNV, and across all three drugs evaluated. In several cases, BBC-corrected ROC–AUC values increased above the diagonal baseline, indicating enhanced predictive performance when clinical variables were integrated. In other settings, performance remained comparable, suggesting that the added clinical information did not uniformly benefit all combinations.

## Discussion

This study systematically evaluated multiple strategies for transferring pre-clinical knowledge to clinical chemotherapy response prediction and reveals a consistent and instructive pattern. Approaches that rely on fixed or indirect forms of transfer—such as biomarker-based feature selection or biologically informed pathway abstractions derived from cell-line experiments—do not yield robust or systematic performance improvements when applied to patient cohorts. Although such strategies are theoretically appealing because they substantially reduce feature dimensionality and mitigate the classical *p* ≫ *n* problem, our results demonstrate that even extensive sets of experimentally validated biomarkers, when combined with flexible machine-learning models, fail to outperform models trained directly on raw omics data. This finding highlights a persistent gap between conclusions drawn from in vitro systems and the biological and clinical reality of human tumours, suggesting that pre-clinical evidence alone may be insufficient and, in some cases, overly optimistic when translated to clinical prediction tasks.

At the same time, our analyses indicate that pre-clinical signals are not entirely absent from clinical data. After basic batch-effect correction, models trained on cell-line data retained certain predictive signal when evaluated on TCGA cohorts, implying partial conservation of drug-response mechanisms across domains. However, direct deployment of pre-clinical deep-learning models remained unstable and highly drug dependent, reinforcing that distributional alignment alone cannot overcome the fundamental biological differences between cell lines and patient tumours. These results clarify that the challenge lies not in the complete absence of transferable information, but in how that information is incorporated and adapted to the clinical context.

Among the evaluated transfer strategies, only methods that allow explicit adaptation to clinical data produced consistent improvements. Fine-tuning pre-trained models on patient cohorts resulted in moderate but reproducible gains and improved stability, supporting its potential as larger and more diverse clinical datasets become available. In parallel, we evaluated a simpler hybrid transfer strategy in which predictions from a pre-clinical model are used as features within clinical machine-learning models. Despite its conceptual simplicity, this approach consistently improved performance relative to direct pre-clinical model application and offered a practical advantage: it enables seamless integration of clinical variables that are inherently unavailable in pre-clinical datasets, such as age, performance status, and tumour grade. Incorporating these patient-level features yielded further gains in a substantial subset of settings, underscoring that clinically relevant determinants of chemotherapy response extend beyond tumour molecular profiles alone.

Several limitations should be considered when interpreting these findings. For practical reasons, fine-tuning and hybrid transfer strategies were primarily evaluated using mixed-cancer test sets, which may introduce a degree of performance overestimation. Because different tumour types vary in baseline drug sensitivity and responder prevalence, models trained and evaluated on pooled cohorts may partially exploit cancer-type differences rather than purely drug-resistance mechanisms. As a result, the reported gains should be interpreted as upper bounds on achievable performance under these settings. In addition, the clinical response labels used in this study are imperfect proxies for true drug sensitivity and are subject to noise arising from imaging-based assessment, tumour heterogeneity, and treatment context, which likely constrains achievable predictive accuracy across all methods.

Despite these limitations, the results of this work provide clear guidance for future efforts in clinical drug response prediction. Strategies that treat pre-clinical knowledge as fixed—whether encoded as biomarkers, pathways, or pretrained representations—are unlikely to generalise reliably without explicit adaptation. In contrast, approaches that allow pre-clinical information to be reweighted, contextualised, or combined with patient-specific data offer a more robust and practically viable path forward. As clinical datasets grow and become more diverse, systematic evaluation of fine-tuning, domain adaptation, and hybrid modelling strategies will be essential. More broadly, these findings emphasise that progress in this field will depend not only on increasingly sophisticated models, but also on careful consideration of domain shift, evaluation design, and the integration of clinical information that captures aspects of drug response beyond tumour molecular state.

## Appendix

### Candidate drug selection criteria

We sought to identify chemotherapeutic agents that are widely used across multiple cancer types and sufficiently represented in both TCGA and GDSC. To this end, we surveyed 172 drugs administered across 30 TCGA cancer types and quantified their suitability using a custom scoring function that accounts for available sample size and diversity of cancer-type usage. For each drug *d*, a composite score was defined as:

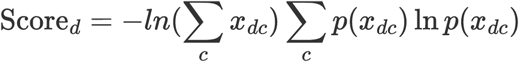

where *x_dc_* denotes the weighted sample size of drug *d* in cancer type *c*, and 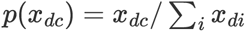. The logarithmic scaling of the total sample size term mitigates the score inflation for drugs with modest absolute counts, while the entropy term favours drugs administered across a broad range of cancer types.

Using this ranking scheme, five drugs emerged as optimal candidates for subsequent analyses (**Table #drug_score#**). Notably, these agents span multiple mechanistic classes, including DNA replication inhibitors and cytoskeletal drugs, and collectively cover hundreds of patients across diverse tumour types. Cisplatin, in particular, achieved the highest score, reflecting both its extensive clinical use and broad cancer-type coverage.

**Table S1.** Drugs selected by customised scoring function.

| Drug | Score | Sample size | Type |
| --- | --- | --- | --- |
| Cisplatin | 10.9 | 346 | DNA replication inhibitor |
| Fluorouracil | 8.48 | 255 | Antimetabolites |
| Gemcitabine | 8.33 | 170 | DNA replication inhibitor |
| Paclitaxel | 9.29 | 211 | Cytoskeletal drug that targets tubulin |

### Literature derived biomarker

**Table S2.** Biomarkers collected from literature. ATC: anaplastic thyroid carcinoma; BTC: Biliary Tract Cancer; BUC: bladder urothelial carcinoma; CCA: cholangiocarcinoma; CDX: Cell line-Derived Xenograft; CL: In-vitro cell line; EAC: esophageal adenocarcinoma; EC: esophageal cancer; GBC: Gallbladder cancer; HCC: hepatocellular carcinoma; HNC: head and neck cancer; MPM: Malignant pleural mesothelioma; OSCC: oral squamous cell carcinoma; PC: pancreatic cancer; PDC: Patient-Derived Cell Cultures; PDS: Patient-Derived Samples (primary patient tumour sample); PDX: Patient-Derived Xenografts; RCT: randomised controlled trial (RCT-H and RCT-M standard for randomised controlled trial based homo sapiens and mice); TNBC: Triple-negative breast cancers; NSCLC: Non-Small Cell Lung Cancer

| Chemotherapy | Biomarker | Cancer and Evidence type |
| --- | --- | --- |
| Cisplatin | ATP7A | Ovarian (CL)<sup>36</sup> |
| Cisplatin | ATP7B | Ovarian (CL)<sup>37</sup> |
| Cisplatin | CTR1/CTR2 | Ovarian (PDS)<sup>38</sup> |
| Cisplatin | ERCC1<sup>39</sup> | Ovarian (CL), colorectal (CL), pancreatic (CL), osteosarcoma (CL), NSCLC (CL) |
| Cisplatin | GSTP1 | Ovarian (CL)<sup>40</sup> |
| Cisplatin | mir-101 | Bladder (CL, CDX)<sup>41</sup>, BUC (PDS, CL)<sup>42</sup> |
| Cisplatin | mir-106a | Cervical (CL)<sup>43</sup>, Ovarian (CL)<sup>44</sup> |
| Cisplatin | mir-106b | Cervical (CL)<sup>43</sup>, Ovarian (CL)<sup>44</sup> |
| Cisplatin | mir-10b | Lung (CL)<sup>45</sup> |
| Cisplatin | mir-124 | NSCLC (CDS, PDS)<sup>46</sup> |
| Cisplatin | mir-125 | GBC (PDS, CL, CDX)<sup>47</sup> |
| Cisplatin | mir-1271 | Colorectal (PDS, CL)<sup>48</sup> |
| Cisplatin | mir-128 | Ovarian (CD, CDX)<sup>49</sup> |
| Cisplatin | mir-1284 | Ovarian (PDS, CL)<sup>50</sup> |
| Cisplatin | mir-129 | GC (PDS, CL)<sup>51</sup> |
| Cisplatin | mir-1293 | Osteosarcoma (CL, CDX)<sup>52</sup> |
| Cisplatin | mir-130a | Ovarian (CL)<sup>53,54</sup> |
| Cisplatin | mir-133b | Lung (CL)<sup>55</sup>, OS (CL)<sup>56</sup> |
| Cisplatin | mir-136 | Ovarian (CL)<sup>57</sup> |
| Cisplatin | mir-137 | Ovarian (CL)<sup>58</sup>, Lung (PDS, CL, CDX)<sup>59</sup> |
| Cisplatin | mir-141 | Ovarian (CL, PDS)<sup>60</sup> |
| Cisplatin | mir-142 | NSCLC (CDS, PDS)<sup>46</sup> |
| Cisplatin | mir-145 | GBC (PDS, CL)<sup>61</sup> |
| Cisplatin | mir-146a | Lung (CL)<sup>62</sup>, NSCLC (CL, PDS)<sup>63</sup> |
| Cisplatin | mir-152 | Ovarian (CL, CDX)<sup>64</sup> |
| Cisplatin | mir-155 | Lung (CL, PDS)<sup>65</sup>, OSCC (CL, PDS)<sup>66</sup>, HCC (CDX, PDS)<sup>67</sup> |
| Cisplatin | mir-15b | Lung (PDS, CL, CDX)<sup>68</sup> |
| Cisplatin | mir-17 | NSCLC (CL, PDS)<sup>69</sup> |
| Cisplatin | mir-181a | NSCLC (CL)<sup>70</sup>, EAC (CL, CDX)<sup>71</sup> |
| Cisplatin | mir-181c | NSCLC (PDS, CL, CDX)<sup>72</sup> |
| Cisplatin | mir-182 | HCC(CL, PDS)<sup>73</sup> |
| Cisplatin | mir-185 | Ovarian (CL, CDX)<sup>64</sup> |
| Cisplatin | mir-186 | Ovarian (CL)<sup>74</sup> |
| Cisplatin | mir-194 | Ovarian (CL)<sup>75</sup> |
| Cisplatin | mir-195 | Gastric (PDS, CL)<sup>76</sup> |
| Cisplatin | mir-196a | HNC (PDS, CL, CDX)<sup>77</sup> |
| Cisplatin | mir-199a | Ovarian (PDS, CL, CDX-Orthotopic)<sup>78</sup>, Breast (CL)<sup>79</sup> |
| Cisplatin | mir-200b | Ovarian (PDS, CL, CDX)<sup>80</sup> |
| Cisplatin | mir-200c | Ovarian (PDS, CL, CDX)<sup>80</sup> |
| Cisplatin | mir-204 | Neuroblastoma (PDS, CL, CDX)<sup>81</sup> |
| Cisplatin | mir-205 | Glioma (CL)<sup>82</sup> |
| Cisplatin | mir-206 | Ovarian (PDS, CL, CDX)<sup>83</sup> |
| Cisplatin | mir-20a | Colorectal (CL, CDX)<sup>84</sup> |
| Cisplatin | mir-21 | Gastric (CL)<sup>85</sup>, NSCLC (PDS)<sup>86</sup> |
| Cisplatin | mir-214 | Osteosarcoma (CL)<sup>87</sup> |
| Cisplatin | mir-216b | Ovarian (CL, CDX, PDS)<sup>88</sup> |
| Cisplatin | mir-218 | Breast (CL, PDS)<sup>89</sup>, Cervical (CL, CDX)<sup>90</sup> |
| Cisplatin | mir-219 | NSCLC (PDS, CL, CDX)<sup>91</sup> |
| Cisplatin | mir-23a | Ovarian (CL)<sup>92</sup> |
| Cisplatin | mir-26a | Gastric (CL)<sup>93</sup> |
| Cisplatin | mir-27a | Lung (PDS, CL, CDX)<sup>94</sup>, Liver (CL)<sup>95</sup> |
| Cisplatin | mir-29 | Ovarian (CL)<sup>96</sup>, NSCLC (CL, CDX, PDS)<sup>97</sup> |
| Cisplatin | mir-29a | NSCLC (PDS, CL)<sup>98</sup> |
| Cisplatin | mir-302b | Breast (CL)<sup>99</sup> |
| Cisplatin | mir-30a | Gastric (CL, PDS)<sup>100</sup> |
| Cisplatin | mir-30b | Lung (PDS, CL, CDX)<sup>101</sup> |
| Cisplatin | mir-30d | ATC (CL, CDX)<sup>102</sup> |
| Cisplatin | mir-324 | Lung (CL)<sup>103</sup> |
| Cisplatin | mir-335 | Ovarian (CL, CDX)<sup>104</sup> |
| Cisplatin | mir-338 | Ovarian (PDS, CL, CDX)<sup>105</sup> |
| Cisplatin | mir-34a | Osteosarcoma (CL)<sup>106</sup> |
| Cisplatin | mir-374a | Ovarian (CL)<sup>53</sup> |
| Cisplatin | mir-375 | Lung (CL)<sup>107</sup> |
| Cisplatin | mir-449a | Gastric (CL)<sup>108</sup> |
| Cisplatin | mir-451 | NSCLC (PDS, CL, CDX)<sup>109</sup> |
| Cisplatin | mir-491 | Gastric (CL)<sup>110</sup> |
| Cisplatin | mir-497 | Ovarian (PDS, CL, CDX)<sup>111</sup> |
| Cisplatin | mir-503 | NSCLC (CL)<sup>112</sup> |
| Cisplatin | mir-556 | NSCLC (CL, CDX)<sup>113</sup> |
| Cisplatin | mir-613 | NSCLC (PDS, CL, CDX)<sup>114</sup> |
| Cisplatin | mir-630 | NSCLC (CL)<sup>70</sup> |
| Cisplatin | mir-664 | Cervical (PDS, CL)<sup>115</sup> |
| Cisplatin | mir-7 | Gastric (CL)<sup>116</sup>, lung adenocarcinoma (PDS)<sup>117</sup>, cervical (CL)<sup>118</sup> |
| Cisplatin | mir-770 | Ovarian (PDS, CL)<sup>119</sup> |
| Cisplatin | mir-9 | NSCLC (CL)<sup>120</sup>, Ovarian (PDC)<sup>121</sup> |
| Cisplatin | mir-92 | NSCLC (CL, PDS)<sup>69</sup> |
| Cisplatin | mir-98 | Ovarian (PDS, CL, CDX)<sup>122</sup> |
| Cisplatin | mir-99a | Gastric (CL)<sup>110</sup> |
| Cisplatin | MRP2 | Colorectal (PDS)[^ Hinoshita E U 2000] |
| Cisplatin | XPA | Testicular (CL)<sup>123</sup> |
| Cisplatin | XPF (ERCC4) | HNSCC (clinical)<sup>124</sup> |
| Fluorouracil | BCRP | Breast (CL)<sup>125</sup> |
| Fluorouracil | mir-106a | Colorectal (PDS, CL)<sup>126</sup> |
| Fluorouracil | mir-106b | CCA (CL, CDX)<sup>127</sup> |
| Fluorouracil | mir-10b | Colorectal (PDS, CL)<sup>128</sup> |
| Fluorouracil | mir-122 | HCC (CL, CDX)<sup>129</sup>, Colorectal (CL, CDX)<sup>130</sup> |
| Fluorouracil | mir-1229 | Gastric (PDS, CL, CDX)<sup>131</sup> |
| Fluorouracil | mir-125b | HCC (CL)<sup>132</sup> |
| Fluorouracil | mir-129 | CRC (PDS, CL, CDX)<sup>133</sup> |
| Fluorouracil | mir-136 | Ovarian (PDS, CL)<sup>134</sup> |
| Fluorouracil | mir-137 | PC (PDS, CL)<sup>135</sup> |
| Fluorouracil | mir-138 | PC (PDS, CL)<sup>136</sup> |
| Fluorouracil | mir-139 | Gastric (PDS, CL, CDX)<sup>137</sup> |
| Fluorouracil | mir-141 | HCC (CL)<sup>138</sup> |
| Fluorouracil | mir-143 | Colorectal (CL)<sup>139</sup> |
| Fluorouracil | mir-145 | Colorectal (PDS, CL, CDX)<sup>140</sup> |
| Fluorouracil | mir-147 | Gastric (PDS, CL)<sup>141</sup> |
| Fluorouracil | mir-17 | Colorectal (CL, PDS)<sup>142</sup> |
| Fluorouracil | mir-181c | Pancreatic (PDS, CL)<sup>143</sup> |
| Fluorouracil | mir-183 | Pancreatic (CL, CDX)<sup>144</sup> |
| Fluorouracil | mir-195 | Colorectal (CL)<sup>145</sup> |
| Fluorouracil | mir-196b | Colorectal (PDS, CL)<sup>146</sup> |
| Fluorouracil | mir-197 | Colorectal (PDS, CL)<sup>147</sup> |
| Fluorouracil | mir-19b | Colon (CL)<sup>148</sup> |
| Fluorouracil | mir-200a | HCC (CL)<sup>149</sup> |
| Fluorouracil | mir-200c | Colorectal (CL)<sup>150</sup> |
| Fluorouracil | mir-203 | Colorectal (CL, CDX)<sup>151</sup> |
| Fluorouracil | mir-204 | Gastric (PDS, CL, CDX)<sup>152</sup> |
| Fluorouracil | mir-206 | Colorectal (CL)<sup>153</sup> |
| Fluorouracil | mir-21 | Pancreatic (CL)<sup>154</sup> |
| Fluorouracil | mir-218 | Colorectal (PDS, CL)<sup>155</sup> |
| Fluorouracil | mir-22 | Colorectal (PDS, CL, CDX)<sup>156</sup> |
| Fluorouracil | mir-221 | Esophageal (CL, PDS, CDX)<sup>157</sup>, Pancreatic (CL)<sup>158</sup> |
| Fluorouracil | mir-223 | Colorectal (clinical, CL)<sup>159</sup> |
| Fluorouracil | mir-23a | Colorectal (PDS, CL)<sup>160</sup> |
| Fluorouracil | mir-27a | HCC (CL)<sup>161</sup> |
| Fluorouracil | mir-302b | HCC (CL)<sup>162</sup> |
| Fluorouracil | mir-320a | Pancreatic (CL)<sup>163</sup> |
| Fluorouracil | mir-329 | Colorectal (PDS, CL)<sup>164</sup> |
| Fluorouracil | mir-338 | EC (PDS, CL, CDX)<sup>165</sup> |
| Fluorouracil | mir-34a | Colorectal (PDS, CL, CDX)<sup>166</sup> |
| Fluorouracil | mir-34b | Endometrial (CL, CDX)<sup>167</sup> |
| Fluorouracil | mir-375 | Colorectal (CL, CDX)<sup>168</sup>, Colorectal (CL)<sup>168</sup> |
| Fluorouracil | mir-4516 | Gastric (CL, CDX)<sup>169</sup> |
| Fluorouracil | mir-494 | Colorectal (CL, CDX)<sup>170</sup> |
| Fluorouracil | mir-503 | HCC (PDS, CL)<sup>171</sup> |
| Fluorouracil | mir-587 | Colorectal (CL, CDX)<sup>172</sup> |
| Fluorouracil | mir-675 | Colorectal (CL)<sup>173</sup> |
| Fluorouracil | mir-761 | Colorectal (PDS, CL)<sup>174</sup> |
| Fluorouracil | mir-96 | Colorectal (CL)<sup>175</sup> |
| Fluorouracil | RRM2 | General<sup>176</sup> |
| Fluorouracil | SLC29A1 | HCC (PDS, CL, CDX)<sup>177</sup> |
| Fluorouracil | Tβ10 | CCA (CL)<sup>178</sup> |
| Fluorouracil | UMPS | HCC (CDX)<sup>179</sup> |
| Gemcitabine | mir-101 | Pancreatic (PDS, CL)<sup>180</sup> |
| Gemcitabine | mir-1243 | Pancreatic (PDS, CL)<sup>181</sup> |
| Gemcitabine | mir-1246 | Pancreatic (PDS, CDX)<sup>182</sup> |
| Gemcitabine | mir-125a | Pancreatic (PDS, CL)<sup>183</sup> |
| Gemcitabine | mir-129 | Bladder (PDS, CL)<sup>184</sup> |
| Gemcitabine | mir-130a | BTC (CL)<sup>185</sup> |
| Gemcitabine | mir-135b | Pancreatic (CL)<sup>186</sup> |
| Gemcitabine | mir-141 | BTC (CL)<sup>187</sup> |
| Gemcitabine | mir-145 | Pancreatic (CL)<sup>188</sup> |
| Gemcitabine | mir-153 | Pancreatic (PDS, CL, CDX)<sup>189</sup> |
| Gemcitabine | mir-155 | Pancreatic (PDS, CL, CDX)<sup>190</sup> |
| Gemcitabine | mir-17 | Lung (CL)<sup>191</sup>, Pancreatic (CL)<sup>192</sup> |
| Gemcitabine | mir-181b | Pancreatic (CL)<sup>193</sup> |
| Gemcitabine | mir-200b | Pancreatic (CL)<sup>194</sup> |
| Gemcitabine | mir-203 | Pancreatic (PDS, CL)[<sup>^</sup>Ren Z G 2016] |
| Gemcitabine | mir-205 | Cholangiocarcinoma (CL)<sup>195</sup> |
| Gemcitabine | mir-20a | Pancreatic (PDS, CL)<sup>196</sup> |
| Gemcitabine | mir-21 | Breast (CL, CDX)<sup>197</sup> |
| Gemcitabine | mir-210 | Pancreatic (PDS, CL, CDX)<sup>198</sup> |
| Gemcitabine | mir-217 | Pancreatic (CL)<sup>199</sup> |
| Gemcitabine | mir-218 | Gallbladder (PDS, CL, CDX)<sup>200</sup> |
| Gemcitabine | mir-221 | Cholangiocarcinoma (CL)<sup>195</sup> |
| Gemcitabine | mir-29 | Cholangiocarcinoma (CL)<sup>195</sup>, Pancreatic (CL)[<sup>^</sup>Nahano H T 2013] |
| Gemcitabine | mir-30a | Pancreatic (PDS, CL, CDX)<sup>201</sup> |
| Gemcitabine | mir-301 | Pancreatic (CL)<sup>194</sup> |
| Gemcitabine | mir-320c | Pancreatic (PDS, CL)<sup>202</sup> |
| Gemcitabine | mir-33a | Pancreatic (CL)<sup>203</sup> |
| Gemcitabine | mir-3656 | Pancreatic (PDS, CL, CDX) |
| Gemcitabine | mir-3662 | Pancreatic (PDS, CL, CDX)<sup>204</sup> |
| Gemcitabine | mir-373 | Pancreatic (CL)<sup>205</sup> |
| Gemcitabine | mir-429 | Pancreatic (CL, CDX)<sup>206</sup> |
| Gemcitabine | mir-497 | Pancreatic (PDS, CL, CDX)<sup>207</sup> |
| Gemcitabine | mir-506 | Pancreatic (CL, CDX)<sup>208</sup> |
| Gemcitabine | mir-509 | Pancreatic (PDS, CL)<sup>181</sup> |
| Gemcitabine | PPP5C | Pancreatic (CL)<sup>209</sup> |
| Gemcitabine | mir-520a | Pancreatic (CL)<sup>209</sup> |
| Gemcitabine | mir-608 | Pancreatic (CL)<sup>210</sup> |
| Gemcitabine | mir-620 | Breast (CL)<sup>211</sup> |
| Gemcitabine | mir-7 | Pancreatic (CL)<sup>212</sup> |
| Gemcitabine | mir-873 | Breast (CL)<sup>213</sup> |
| Gemcitabine | mir-93 | Pancreatic (CL)<sup>214</sup> |
| Gemcitabine | mir-9 | Breast (PDS, CL, CDX)<sup>215</sup> |
| Gemcitabine | SNHG14 (expression) | Pancreatic (CL)<sup>216</sup> |
| Gemcitabine | NEAT1 | Pancreatic (CL, CDX)<sup>208</sup> |
| Paclitaxel | AKT/ERK (pathway) | Gastric (CL)<sup>217</sup> |
| Paclitaxel | EGFR (CNV) | NSCLC (RCT-H)<sup>218</sup> |
| Paclitaxel | HER2 | Breast (CL)<sup>219</sup> |
| Paclitaxel | MAPT | Gastric (PDC)<sup>220</sup> |
| Paclitaxel | mir-100 | Breast (CL)<sup>221</sup> |
| Paclitaxel | mir-105 | Ovarian (PDS, CL, CDX)<sup>222</sup> |
| Paclitaxel | mir-106a | Ovarian (CL)<sup>223</sup> |
| Paclitaxel | mir-107 | Breast (PDS, CL)<sup>224</sup> |
| Paclitaxel | mir-1204 | Nasopharyngeal carcinoma(CL, CDX)<sup>225</sup> |
| Paclitaxel | mir-125a | Cervical (CL)<sup>226</sup> |
| Paclitaxel | mir-125b | Breast (CL)<sup>227</sup> |
| Paclitaxel | mir-134 | Ovarian (PDS, CL)<sup>228</sup> |
| Paclitaxel | mir-135a | NSCLC, Breast, Ovarian , Prostate (CL, CDX)<sup>229</sup> |
| Paclitaxel | mir-142 | Breast (CL, CDX)<sup>230</sup> |
| Paclitaxel | mir-145 | Ovarian (PDS, CL, CDX)<sup>231</sup> |
| Paclitaxel | mir-148 | Prostate (CL)<sup>232</sup> |
| Paclitaxel | mir-149 | Ovarian (CL)<sup>233</sup> |
| Paclitaxel | mir-150 | Ovarian (PDS, CL)<sup>234</sup> |
| Paclitaxel | mir-152 | Breast (PDS, CL)<sup>235</sup> |
| Paclitaxel | mir-155 | Breast (CL)<sup>236</sup> |
| Paclitaxel | mir-16 | NSCLC (CL)<sup>237</sup> |
| Paclitaxel | mir-17 | NSCLC (CL)<sup>237</sup> |
| Paclitaxel | mir-181c | Ovarian (PDS, CL, CDX)<sup>238</sup> |
| Paclitaxel | mir-186 | Ovarian (CL)<sup>239</sup> , Breast (PDS, CL)<sup>240</sup> |
| Paclitaxel | mir-186 | NSCLC (PDS, CL, CDX)<sup>241</sup> |
| Paclitaxel | mir-18a | Breast (PDS, CL)<sup>242</sup> |
| Paclitaxel | mir-194 | Ovarian (CL)<sup>243</sup> |
| Paclitaxel | mir-197 | Ovarian (CL)<sup>244</sup> |
| Paclitaxel | mir-199a | Prostate (PDS, CL, CDX)<sup>245</sup> |
| Paclitaxel | mir-200c | Ovarian (CL)<sup>246</sup> , Breast (CL)<sup>247</sup> |
| Paclitaxel | mir-203 | Colorectal (PDS, CL)<sup>248</sup> |
| Paclitaxel | mir-21 | Breast (PDS, CL)<sup>249</sup> , Ovarian (CL)<sup>250</sup> |
| Paclitaxel | mir-22 | Colon (CL)<sup>251</sup> |
| Paclitaxel | mir-24 | Breast (PDS, CL, CDX)<sup>252</sup> |
| Paclitaxel | mir-27a | Overian (CL)<sup>253</sup> |
| Paclitaxel | mir-29c | Nasopharyngeal carcinoma(CL, CDX)<sup>254</sup> |
| Paclitaxel | mir-30a | NSCLC (PDS, CL, CDX)<sup>255</sup> |
| Paclitaxel | mir-31 | Ovarian (CL)<sup>256</sup> |
| Paclitaxel | mir-34a | Prostate (CL)<sup>257</sup> |
| Paclitaxel | mir-375 | Cervical (PDS, CL, CDX)<sup>258</sup> |
| Paclitaxel | mir-421 | NSCLC (CL)<sup>259</sup> |
| Paclitaxel | mir-433 | Ovarian (CL)<sup>260</sup> |
| Paclitaxel | mir-451 | Breast (CL, CDX)<sup>261</sup> |
| Paclitaxel | mir-490 | Ovarian (CL)<sup>262</sup> |
| Paclitaxel | mir-520h | Breast (PDS, CL)<sup>263</sup> |
| Paclitaxel | mir-522 | Ovarian (CL)<sup>264</sup> |
| Paclitaxel | mir-591 | Ovarian (CL)<sup>223</sup> |
| Paclitaxel | mir-621 | Breast (clinical, CL)<sup>265</sup> |
| Paclitaxel | mir-634 | Nasopharyngeal carcinoma (CL, CDX)<sup>266</sup> |
| Paclitaxel | mir-7 | NSCLC (CL)<sup>267</sup>, Breast (CL)<sup>268</sup>, Breast (PDS, CL)<sup>240</sup> |
| Paclitaxel | mir-98 | Endometrial (PDS, CL, CDX)<sup>269</sup> |
| Paclitaxel | P-gp/MDR1 | Pan cancer (RCT-M)<sup>270</sup> |
| Paclitaxel | TP53 (status) | Ovarian (clinical)<sup>271</sup> |
| Paclitaxel | UCA1 | Ovarian (CL)<sup>272</sup> |
| Paclitaxel | βIII-tubulin | Gastric (CL)<sup>273</sup> |

### Sample distribution in drug cohort

**Figure S1.**
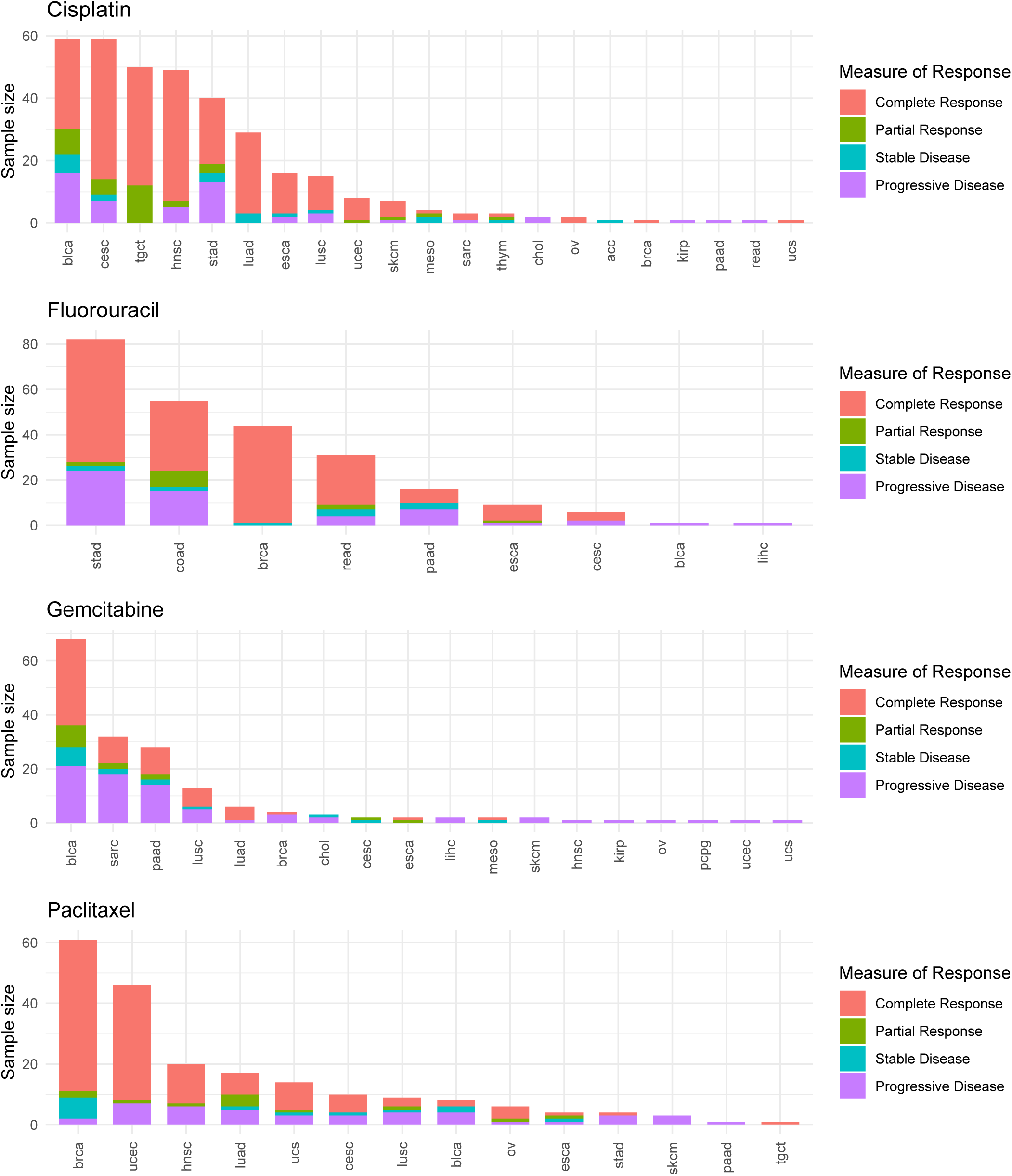
Sample distribution in each of the four drug cohort.

### Biomarker performance in specific types of cancer

**Figure S2.**
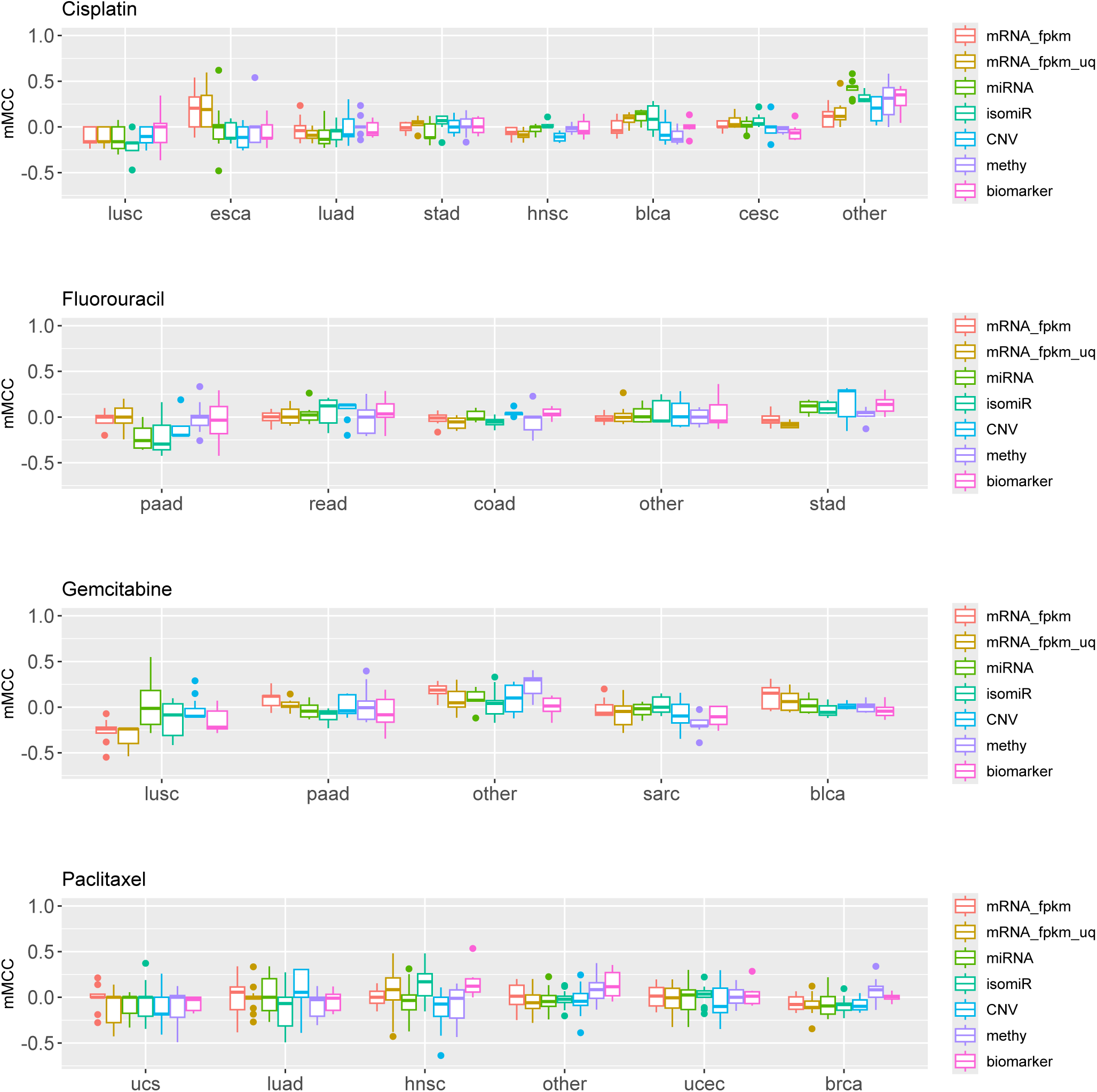
performance of mix-cancer models trained on all omics types and biomarkers. The x-axis indicates the cancer type of the test set, and the y-axis indicates the median MCC distribution of the 16 algorithms (8 for the biomarker-based model) from 5 repetitions of evaluation with different random seeds.

